# Influence of the mutation load on the genomic composition of hybrids between outcrossing and self-fertilizing species

**DOI:** 10.1101/2022.12.12.520111

**Authors:** Fréderic Fyon, Waldir M. Berbel-Filho

## Abstract

Hybridization is a natural process whereby two diverging evolutionary lineages reproduce and create offspring of mixed ancestry. Differences in mating systems (e.g., self-fertilization and outcrossing) are expected to affect the direction and extent of hybridization and introgression in hybrid zones. Among other factors, selfers and outcrossers are expected to differ in their mutation loads. This has been studied both theoretically and empirically; however, conflicting predictions have been made on the effects mutation loads of parental species with different mating systems can have on the genomic composition of hybrids. Here we develop a multi-locus, selective model to study how the different mutation load built up in selfers and outcrossers as a result of selective interference and homozygosity impact the long-term genetic composition of hybrid populations. Notably, our results emphasize that genes from the parental population with lesser mutation load get rapidly over-represented in hybrid genomes, regardless of the hybrids own mating system. When recombination tends to be more important than mutation, outcrossers’ genomes tend to be of higher quality and prevail. When recombination is small, however, selfers’ genomes may reach higher quality than outcrossers’ genomes and prevail. Taken together these results provide concrete insights into one of the multiple factors influencing hybrid genome composition and introgression patterns in hybrid zones with species containing species with different mating systems.

## 1. Introduction

Hybrid individuals are defined as progeny of mixed ancestry; they bear genes inherited from divergent evolutionary lineages. Introgressive hybridization (i.e., formation of hybrids with backcrossing) functions as bridges between divergent lineages, allowing gene flow and shared evolutionary history. Identifying the processes shaping the evolutionary fate of hybrid populations is a fundamental task for evolutionary biology, as well as for conservation, and wildlife management.

The possibility of hybridization happening, along with its ecological and genomic consequences, depends, among other factors, on the mating systems and reproductive biology of the parental species and their hybrids. Most plants and many animals are simultaneous hermaphrodites (Jarne and Auld 2006; The Tree of Sex Consortium 2014). In these species, self-fertilization (selfing) – offspring produced by the fusion of gametes stemming from different meiosis in the same individual – is a possibility. Selfing contrasts with outcrossing as in the latter the progeny is produced by the fusion of gametes proceeding from different meiosis in different individuals (hermaphrodites or not). Mating systems are usually defined in three main categories according to the selfing rate: obligate outcrossing (selfing rate ≤ 20%), predominantly selfing (selfing rate > 80%), and mixed-mating (selfing rate > 20% and ≤ 80%) (Shimizu and Tsuchimatsu 2015). It is only recently that attention has been brought on the effects that different mating systems have on the occurrence of hybridization and on the genomic composition of hybrid populations (Hu 2015; Kim *et al*. 2018; Pickup *et al*. 2019).

Hybrid zones are regions of contact between divergent species/lineages that often produce individuals of admixed ancestry through interspecific mating. Hybrid zones between species with different mating systems (e.g., predominantly selfing and obligate outcrossing) are commonly reported in plants (Ruhsam *et al*. 2011; Pickup *et al*. 2019; Ostevik *et al*. 2021), but examples are scarce in animals (Berbel-Filho *et al*. 2021). Some predictions on how reproductive systems (Brandvain and Haig 2005) and reproductive timing (Martin and Willis 2007; Berbel-Filho *et al*. 2021), along with other ecological factors (Busch *et al*. 2022), can influence the occurrence and outcome of hybridization in these cases have been extensively provided. On the other hand, few studies have tried to address how mating systems differences influence (i) the quality of each parental genomes inherited by first-generation hybrids (F1), and (ii) how the genetic composition (defined in terms of genes’ species-of-origin) of hybrids evolves in the long-term as a result (Hu 2015; Kim *et al*. 2018; Pickup *et al*. 2019). Disparate, sometimes contradictory predictions on these points have been made; the present work was designed to disentangle these and provide a unified framework for the influence of mutation load on hybrid ancestry.

Predictions concerning hybrids’ genetic composition and evolution have focused on mutation load. They rely on the idea that species with different mating systems are expected to harbor different mutation loads (Arunkumar *et al*. 2015; Pickup *et al*. 2019). High homozygosity caused by long-term selfing means that recombination is – most of the time – inefficient at shuffling genetic combinations, which results in higher rates of hitch-hiking, background selection, and selective interference (Wright *et al*. 2013). That is, selection is expected to be less efficient in selfing populations, which are thus predicted to have more slightly deleterious alleles become fixed by random genetic drift (Charlesworth *et al*. 1993). This relates to the classic argument that the absence of effective recombination in asexual and self-fertilizing species should lead them to extinction due to mutational meltdown (Gabriel *et al*. 1993; Lynch *et al*. 1993; Lynch *et al*. 1995). Altogether, this predicts that genomes of self-fertilizing species tend to be of worse quality. On the other hand, however, highly deleterious recessive alleles are expected to be purged by selection in selfers as they are “exposed” to selection by high homozygosity, while in outcrossers they can be maintained at low frequencies as they are partly hidden from selection in heterozygotes (Wang *et al*. 1999). In other words, here selection is predicted to be more efficient in self-fertilizing species, resulting in genomes of better quality in selfers. Overall, selfers’ genomes are expected to harbor more fixed slightly deleterious, codominant mutations and less strongly deleterious, recessive mutations (Arunkumar *et al*. 2015). How are these two alternative mutation loads expected to interact and determine genetic composition and evolution of hybrid populations?

The answer to this question is likely to depend on several parameters of the selection processes in parental species, but also on the hybrid mating system. Highly deleterious, recessive mutations coming from outcrossing ancestors may rapidly be purged by selection if the hybrids self-fertilize (as these mutations rapidly become homozygous and visible to selection), taking together any linked genes. This process is likely to limit the introgression of outcrossers’ genomes into selfers’ (Pickup *et al*. 2019). On the other hand, the many fixed slightly deleterious, codominant mutations coming from the selfing ancestors should be purged out in an outcrossing hybrid, this time restricting introgression of self-fertilizers’ genomes into outcrossers’ (Pickup *et al*. 2019). Under this argument, everything happens as if each parental genome was “adapted” to the mating system it has evolved in and tends to fare worse in alternative mating systems. This raises the question, does the mating system of the hybrid population have a critical influence on the evolution of hybrid genome composition, or is this simply determined by the respective quality of the parental populations’ genomes?

Here, we aim at answering these questions by building a multi-locus model allowing for selection, recombination, mutation, and drift to interplay and modify simple genomes. Running numerical simulations for a diversity of scenarios, we were able to study the effects of selective interference and homozygosity on (1) the relative mutation loads of self-fertilizing and outcrossing populations, and (2) the long-term genetic composition of hybrid populations deriving from parental populations with different mating systems.

First, our work allows to confirm some of the classic arguments on mutation load in self-fertilizing species. As another theoretical study showed recently (Sianta *et al*. 2022), how mutation loads in self-fertilizing and outcrossing populations compare with one another strongly depends on the recombination rate between selected loci. At high recombination, outcrossers appear to be more efficient at limiting mutation load, as frequent reshuffling of genes significantly reduce selective interference and fixation of deleterious mutations. However, at low recombination regime, outcrossers suffer from selective interference just as much as selfers. The only difference between the two then reside in the homozygosity of selfers that purge strongly deleterious recessive mutations, resulting in them having a lesser mutation load relative to outcrossers.

Second, we were able to determine the parameters that have a notable influence on the genetic composition and evolution of the hybrid population. Interestingly, we show that the mating system of the hybrid population does not largely influence the outcome of the simulations. Hybrid genetic composition in the long term appears to be primarily dictated by the relative mutation load of the parental species: genomes of worse quality get preferentially eliminated. This result provides a new key to understand empirical data related to hybrid genome composition and patterns of introgression in hybrid zones between selfer and outcrosser species. Depending on genomic (recombination rates, mutation rates), ecological (intensity of selection), and demographical (population size, bottlenecks, colonization events) parameters, one can now predict how mutation load may have built in the parental species, and how much this could have contributed to hybrid genome composition.

## 2. Methods

We used SLiM3 (Haller and Messer 2019) software to code and run the model. SLiM3 has been used in a diversity of population genetics studies as it provides a resource-effective and flexible computing framework. Because the model is of a stochastic nature, each simulation is repeated 100 times, and mean values as well as standard deviations are calculated from these 100 iterations. The code used here is available on the following GitHub repository: https://github.com/FredericFyon/Mating-systems.

### 2.1. Theoretical populations

Our aim here is to follow the genetic alleles at different loci in populations with different mating systems, along with their hybrids. To do so, we always model three populations: (1) an exclusively or mostly self-incompatible (SI) population that reproduces via obligate outcrossing (outcrossers); (2) a self-compatible (SC) population that reproduces exclusively or mostly by means of self-fertilization (selfers); (3) a population born by hybridization of the two previous ones. Populations do not interact with each other (except at the one generation where hybridization happens) and do not compete. Each population size is set at *N_pop_.* In the following, we call *σ_i_* the rate of self-fertilization of a given population *i*. We refer to the self-fertilization rate of the outcrossing, self-fertilizing, and hybrid populations as *σ_o_ ≤ 0.1*, *σ_s_ ≥ 0.9* and *σ_h_* respectively.

### 2.2. Genetic architecture

The genome of all individuals in the three populations consists of *N_loc_* loci, equally distributed along a chromosome. We note *r* the recombination rate between two adjacent loci. We consider here an infinite-allele model: each mutation occurring is unique, all loci potentially have an infinite number of alleles. In this model we only consider deleterious mutations and note *s* the detrimental effect of a particular mutation on the fitness of the host individual.

### 2.3. Life Cycle

At each generation, selection is modelled by choosing the parents of every offspring born at this generation. To do that, we first determine if an offspring was produced by self-fertilization or not. With probability σ, it is produced by self-fertilization. In that case, we randomly draw one parent from the distribution of individuals of the same population at the previous generation. In contrast, with probability 1 - σ, the offspring is producing by outcrossing: we randomly draw two parents. Drawn parents are then accepted with a probability equal to their fitness *w_i_*. We assume fitness to be purely multiplicative among loci (we assume no epistasis): *w_i_* = Π*_j_ w_i,j_*, with *w_i,j_* being the contribution of locus j [1, *N_loc_*] to the fitness of individual *i*. *w_i,j_* depends on (1) the detrimental effect 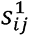 and 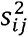 of the alleles present on the two homologous chromosomes at locus *j* of individual *i*, and on (2) the dominance coefficients 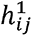 and 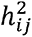 of these two alleles. Following standard population genetics design, if the alleles are not identical-by-descent (that is, if the individual is heterozygote), *w_i,j_* is calculated as:

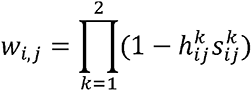

If they are identical by-descent (homozygote), then *w_i,j_* simply is: *w_i,j_* = 1 − *s_i,j_* (with 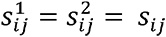).

Following empirical data suggesting that the dominance of a deleterious mutation is negatively correlated with its detrimental effect (Simmons and Crow 1977; Phadnis and Fry 2005; Agrawal and Whitlock 2011), we assume in this model a negative exponential relationship between *s* and *h*:

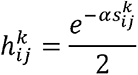

with being an arbitrarily chosen parameter that allows to modulate this relationship. Here, we will always assume that for simplicity. This allows to approximately match empirical data of mean dominance of mildly deleterious mutations (*s = 0.1*) being around 0.25 (Manna *et al*. 2011; however see Simmons and Crow 1977; García-Dorado & Caballero 2000; for example, for other estimates). The exponential function allows for a long-tailed asymptotic distribution of dominance effects, letting highly deleterious mutations to have a very low but strictly positive dominance. Also, the function means that slightly deleterious mutations approach codominance. This allows us to incorporate in the model deleterious mutations in the whole range of dominance values from almost complete recessivity to almost codominance. Overall, this allows the whole selective process to be determined by only one parameter: the distribution of detrimental effects of deleterious mutations.

In addition to selection, the life cycle includes recombination and mutation. Recombination happens at a rate *r* being any two adjacent loci. A recombination event exchanges the composition of the two homologous chromosomes in an individual from the recombination point to the downstream end on the chromosomes. Of course, multiple recombination events can take place during any meiotic event, leading to potentially mosaic inherited chromosomes.

Mutations happen with probability *μ* per locus per chromosome per individual. When a mutation occurs, we draw its detrimental effect from a negative exponential distribution of mean *1/λ*. This means that there is a vast majority of slightly deleterious, codominant mutations arising, along with some mildly deleterious, recessive mutations and very few highly deleterious, strongly recessive mutations.

### 2.4. Hybridization

At first, we consider only the parental populations of outcrossers and selfers. These reproduce within themselves for a certain number of generations *t_1_*. Then, a hybridization event happens. Hybridization is modelled by created a third, hybrid population. This population is created by drawing for each individual one parent from each parental population. As a result, at *t_1_ + 1*, every individual of the hybrid population bears one chromosome from one parental population, and the other from the other parental population. After that initial hybridization event, we consider that there is no additional gene flow from the parental populations (as backcrossing would blur the sole effect of mutation load). We let the hybrid population reproduce within itself and evolve in the absence of additional mutation. This allows us to determine the evolutionary fate of the parental genomes within the hybrid population.

### 2.5. Parameter values

Unless specified otherwise, simulations throughout this work have been run with the following parameter values: Npop = 500, Nloc = 100, λ = 10, σ_o_ = 0 (purely outcrossing population) and σ_s_ = 1 (purely self-fertilizing population). We discuss later the influence of these parameters and provide examples with some other values in the Supplementary Material.

## 3. Results

### 3.1. Mutation Load in Outcrossers and Selfers

We present in the Supplementary Material some results of how the mutation load builds up in outcrossing and self-fertilizing populations. We only rapidly present these here as they are already known patterns (Sianta *et al*. 2022).

Following classic population genetics arguments, our model shows that outcrossers’ genomes present higher heterozygosity, more deleterious mutations, and lower dominance coefficients than selfers’ genomes.

Our results also emphasize that the relative rates of fixation of deleterious mutations in outcrossers and selfers highly depend on mutation and recombination rates. Higher mutation rates mean more fixations. Interestingly, we see that when recombination rate is large, selfers accumulate more fixed deleterious mutations, as they suffer from selective interference: high rates of homozygosity mean recombination is mostly inefficient at reshuffling genetic associations. However, when recombination rates are small, outcrossers suffer from selective interference just as much as selfers. Consequently, outcrossers may accumulate more deleterious mutations than selfers, as they suffer from a reduction of selection efficacy associated with the detrimental effect of mutations being partially hidden in heterozygotes.

Overall, this allows us to determine areas of the parameter space where outcrossers’ genomes are of better quality (high recombination rates), and areas of the parameter space where selfers’ genomes are of better quality (low recombination rates). Parting from here, we can investigate how these genomes fare when put together in hybrids.

### 3.2. Genetic Composition of Hybrids between Outcrossers and Selfers

In the following, we look at the proportions of genes in the hybrid population coming from each of the parental populations. On Fig. 1, we illustrate the genomic composition of hybrid populations, considering a case where *μ = 10^-5^* and where the parental outcrossing and self-fertilizing populations have diverged during 10,000 generations prior to hybridization. We associate these patterns with the mutation load built-up in the parental populations.

**Fig. 1:**
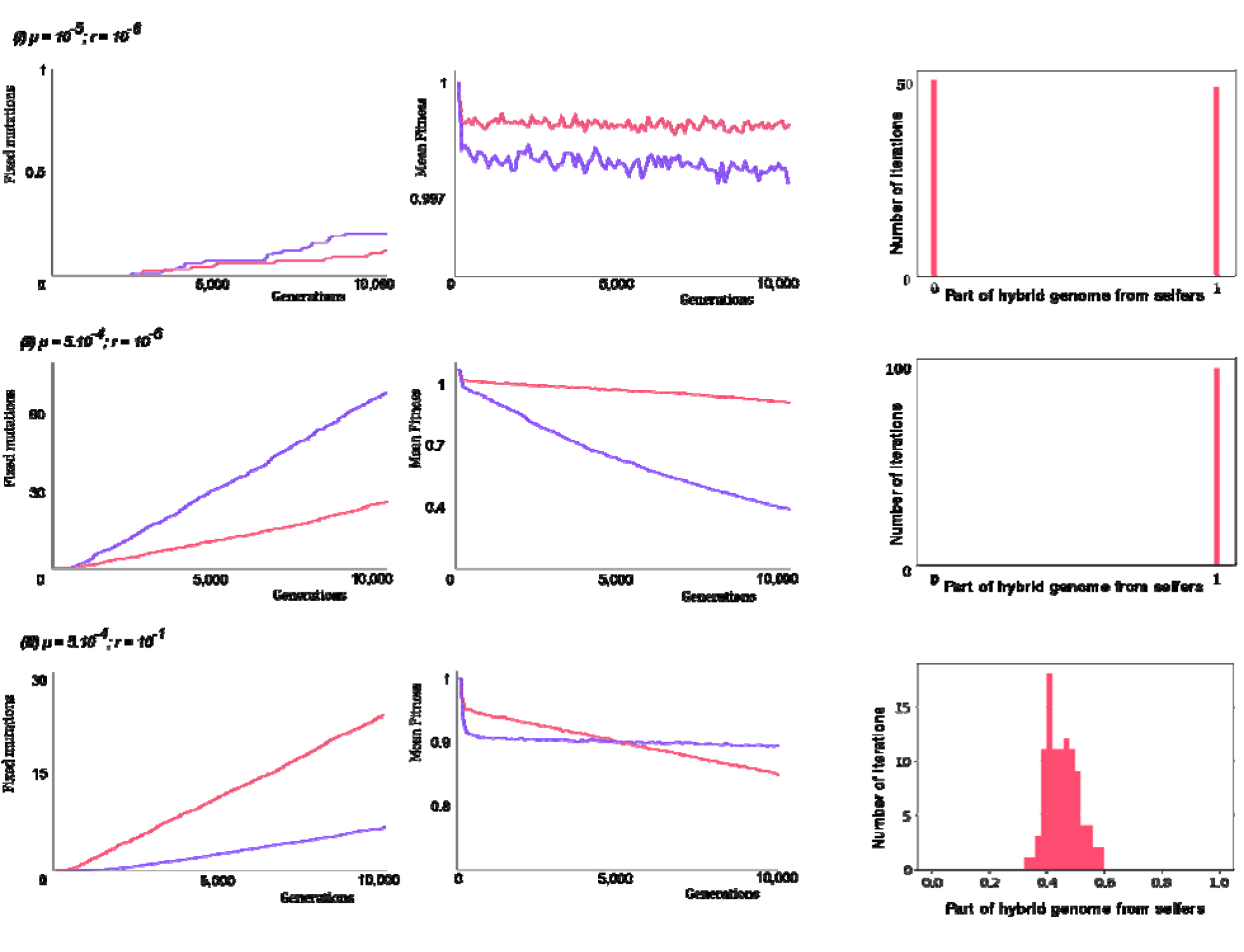
Consequences of the accumulation of mutation load in the parental populations on the genomic composition of hybrids. Here, an outcrossing (purple) and a self-fertilizing (red) populations evolve for 10,000 generations. First column shows the number of fixed deleterious mutations in their respective genomes across different mutation and recombination rates (i, ii, iii). Second column shows the respective population mean fitnesses. Simulations are repeated 100 times: plain lines account for average values, while shaded areas display the standard deviation. After 10,000 generations, the parental populations hybridize, and we look at the genomic composition of the hybrid population after 20,000 additional generations. Column 3 displays histograms of the part of hybrid genome coming from the self-fertilizing population. Simulations are run for three different mutation and recombination rates:(i) µ = 10^-5^ and r = 10^-6^ (first row), (ii) µ = 5.10^-4^ and r = 10^-6^ (second row), and (iii) µ = 5.10^-4^ and r = 10^-1^ (third row).

Figure 1 exemplifies three different scenarios. In scenario (i), the mutation rate and time left for mutation accumulation are so small that mutation load barely builds up. Outcrossing and self-fertilizing populations accumulate similar amounts of deleterious mutations. As a result, they do not differ much in population mean fitness. The outcrossing population has lower fitness due to higher frequencies of highly deleterious, strongly recessive mutations, but the difference is marginal. As a result, in the hybrid population around half of the time the genes of the outcrossing parents prevail, half the time the genes of the self-fertilizing parents prevail. We see that there is no case of mixture of genes of the two populations. This is because the recombination rate is too low here. Eventually, one haplotype goes to fixation, and because the recombination rate is so small this haplotype is one chromosome of either one of the parental populations. The two populations having similar mean fitnesses; in fact, they both have many chromosomes without any deleterious mutations. Most probably, one of these “optimal” chromosomes goes to fixation in the hybrid population, and there is almost equal probability that the “optimal” haplotype that goes to fixation is from one or the other parental populations.

The results of scenario (i) strongly rely on the fact that deleterious mutations are rare, so that many genomes are completely free of them or harbor very few of them. In the following scenarios, we consider alternative cases where there are many detrimental mutations around, either fixed or segregating. Here, we use mutation rate values that are not realistic but allow us to simulate a scenario with many mutations emerge, which is arguably more realistic than the scenario (i) where most genomes are “perfect”.

In scenario (ii) there is significant accumulation of deleterious mutations. Because the recombination rate is still low, both the outcrossing and the self-fertilizing populations suffer from selective interference. Heterozygosity in the outcrossing populations helps deleterious mutations go to fixation; as a result, we see in this scenario that the outcrossing population accumulates more fixed deleterious mutations than the self-fertilizing population. This translates in the outcrossing population having a much lower population mean fitness after 10,000 generations than the outcrossing population. In this case, we see that the genes from the self-fertilizing population always prevail in hybrids. Again, because recombination rate is small, the haplotype that goes to fixation is an entire chromosome of one of the parental populations, here always the self-fertilizing population since its genomes are of much better quality.

In scenario (iii), high recombination rates mean that the outcrossing population suffer much less from selective interference than the self-fertilizing population. Therefore, it accumulates less fixed deleterious mutations, and have a somewhat better fitness after 10,000 generations than the self-fertilizing population. In this case, we see that on average the genes from the outcrossing population prevails. However, we see that hybrid genomes are a mixture of genes from the two parental populations. This is because here high recombination rates means that a mosaic haplotype goes to fixation. Because the genomes of the outcrossing population are of better quality here, this mosaic tends to contain more genes from the outcrossing parents than genes from the self-fertilizing parent.

### 3.3. Effects of hybrid mating system on hybrid genome composition

On Fig. 1, the hybrid population always outcross to reproduce. To gain insights into how the mating system of the hybrid population impacts its genome composition, we show on Fig. 2 histograms of the part of the genome of the hybrid populations that stems from the self-fertilizing parental populations for different values of *r*, *µ*, but also *σ_h_*, the probability of hybrids to reproduce via self-fertilization.

**Fig. 2:**
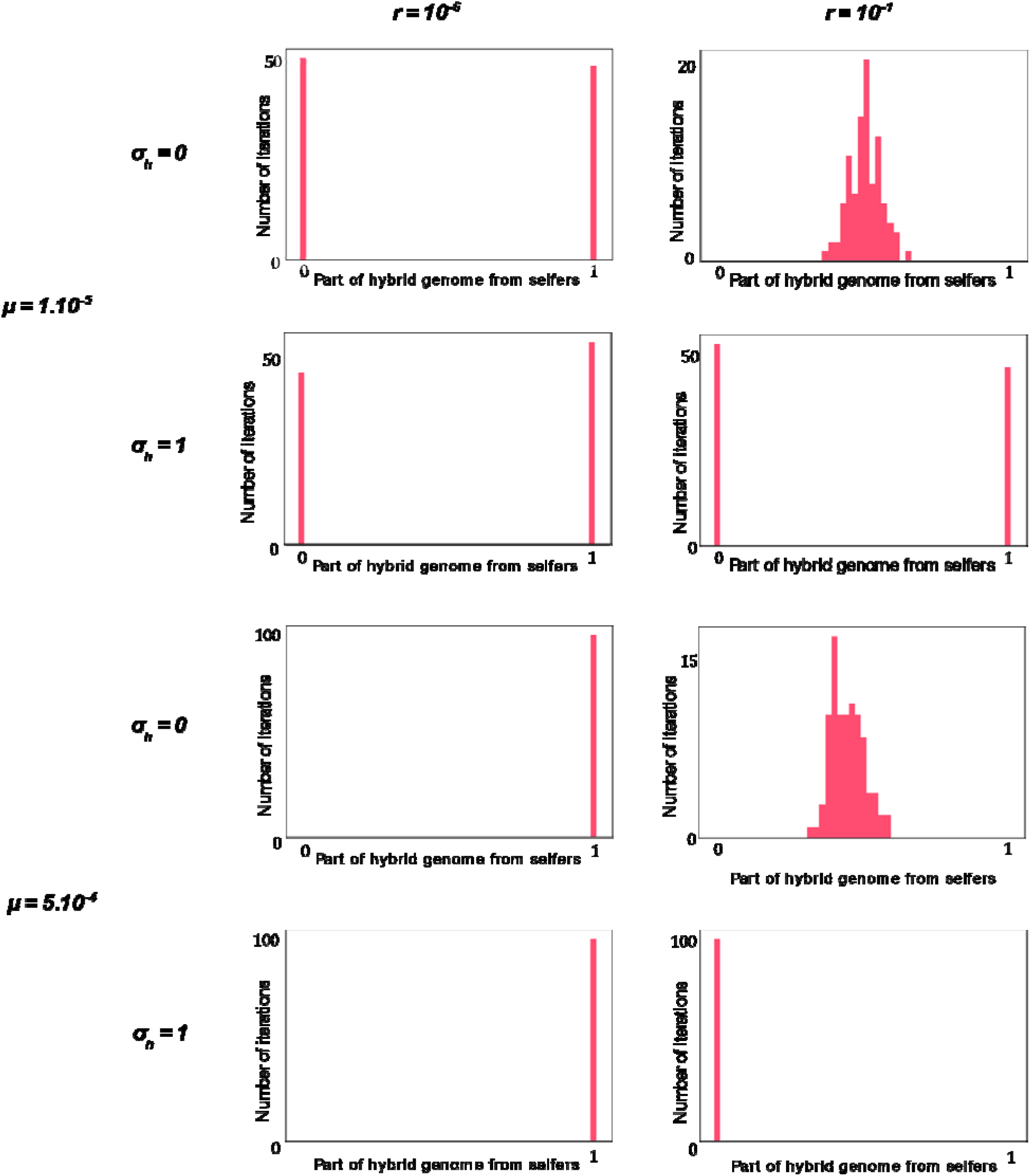
Hybrid genome composition depends on µ, r and σ_h_. We illustrate here histograms of the part of the hybrid genomes that were inherited from the self-fertilizing parental population. Results are displayed for various recombination rates (first column: r = 10^-6^, second column: 10^-1^), mutation rates (top four rows: µ = 10^-5^, bottom four rows: µ = 5.10^-4^), and rates at which the hybrid population reproduces by self-fertilization (first and third rows: σ_h_ = 0, second and fourth rows: σ_h_ = 1).

The mating system of the hybrid seems to have only a marginal effect on the genomic composition of the hybrids. Regardless whether hybrids self-fertilize or outcross, it does not change which genes tend to prevail. When the genomes of the two parental populations are of similar quality (*µ = 1.10^-5^*), the probability for each type of genome to prevail remains around one half whether the hybrids self-fertilize or outcross. When the genomes of the self-fertilizing parents tend to prevail (*µ = 1.10^-4^, r = 10^-6^*), they prevail also in both cases. Finally, when the genomes of the outcrossing parents tend to prevail (*µ = 1.10^-4^, r = 10^-1^*), hybrid genomes are biased towards outcrossing parents’ heredity again in both cases.

The effect of the mating system mostly appears when *r = 10^-1^*. In that case, there is a fundamental difference depending on how the hybrids reproduce. If the hybrids outcross, then their genomes are frequently recombined. As evoked earlier, this means that hybrids genomes become a mixture of the two parental populations, instead of being only composed of genes from one parental populations. When *µ = 5.10^-4^*, this means that the genomes of the hybrids will be composed of a mixture of genes biased towards outcrossing heredity instead of being composed exclusively of genes from the outcrossing population.

Overall, we see that when there are many mutations around, the genomic composition of hybrids between species with different mating systems mostly depends on the recombination rate between loci. If recombination is low, outcrossers accumulate more mutations than selfers and their genome tend to disappear in hybrids. In contrast, if recombination rate is high, selfers’ genomes accumulate more deleterious mutations than outcrossers and their genes tend to disappear in hybrids. Overall, the hybrid mating system seems to have a marginal impact: the hybrid genomic composition seems primarily directed by parental populations’ mutation load and genome quality.

Fig. 2 was drawn for complete self-fertilization (*σ_o_ = 1* and *σ_h_ = 1*). However, in nature, self-fertilization is often associated with low rates of outcrossing (Jarne and Auld 2006). To take that into account, we show on Fig. 3 the effect of assuming self-fertilization rates of only 90%. We see that most of the time, low rates of outcrossing do not change the average ancestry in hybrids (around 50% of hybrid genome coming come the self-fertilizing parent when *µ = 10^-5^*, only the self-fertilizing parents’ genes transmitted to hybrids when *µ = 10^-5^* and *r = 10^-6^*). However, patterns reverse in the case where *µ = 10^-5^* and *r = 10^-1^*. For these parameter values, only outcrossing parents’ genes are inherited in hybrids when *σ_o_ = σ_h_ = 1*, while a majority of self-fertilizing parents’ genes are transmitted when *σ_o_ = σ_h_ = 0.9*. We saw on Fig. 1 that for these parameter values, outcrossing species’ mean fitness just slightly overcome selfing species’ mean fitness. When the self-fertilization rate of the selfing species is increased, it accumulates less deleterious mutations, enough for it to gain better fitness and its genes to be preferentially transmitted in hybrids. Overall, we see again that ancestry patterns will depend on the accumulated mutation load in the parental species, which will depend on parameters such as *µ* and *r*, as seen before, but also the exact reproductive modes of each of the parental species.

**Fig. 3:**
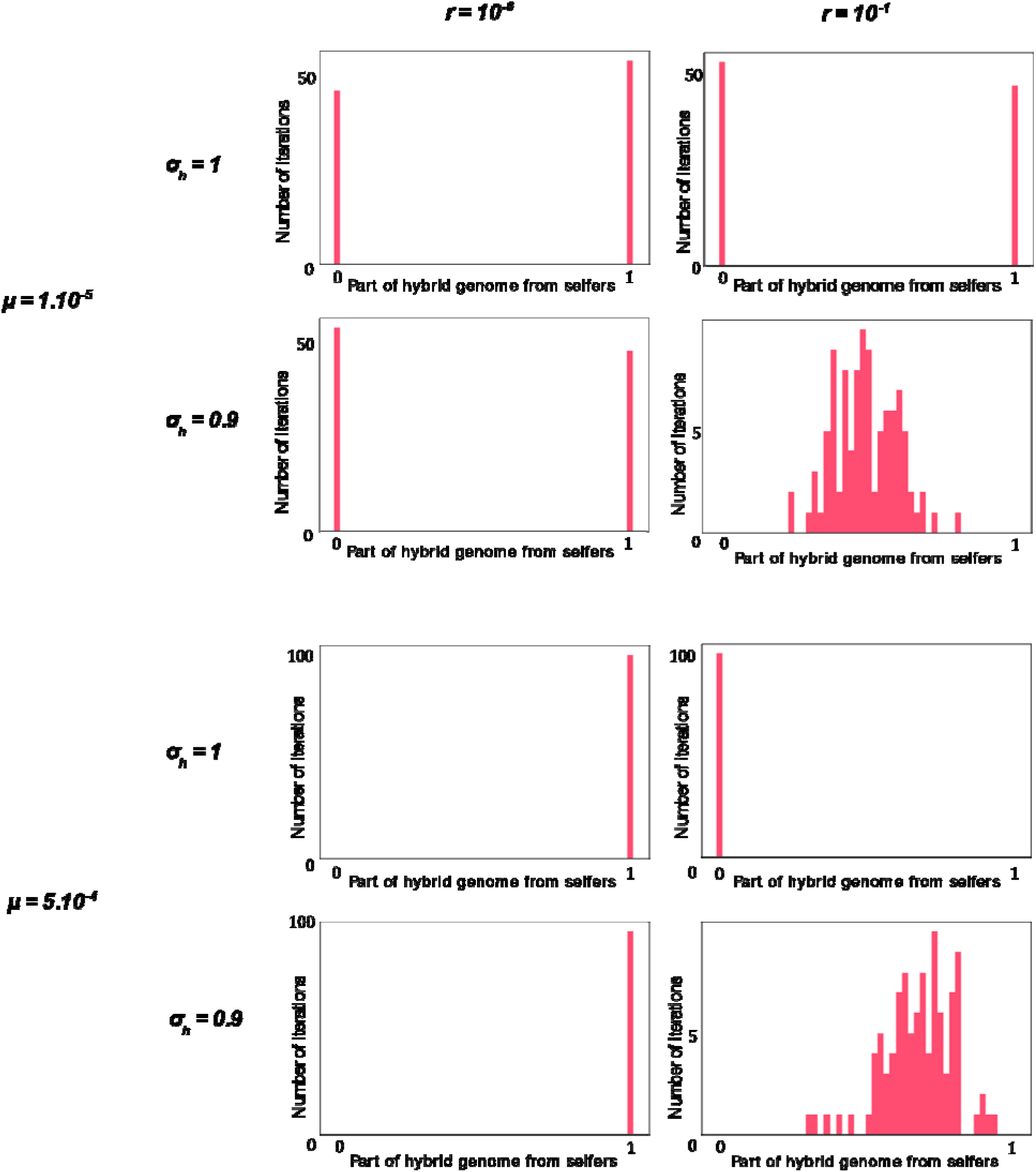
Hybrid genome composition also depends on the exact reproductive modes. Same values of r and µ are used as for Fig. 2. We now compare σ_o_ = σ_h_ = 1 (first and third rows), with σ_o_ = σ_h_ = 0.9. Again, histograms are shown for 100 iterations, with the x-axis representing the part of the hybrid genome that, 10,000 generations after hybridization, was inherited from the self-fertilizing parents.

**Fig. 4:**
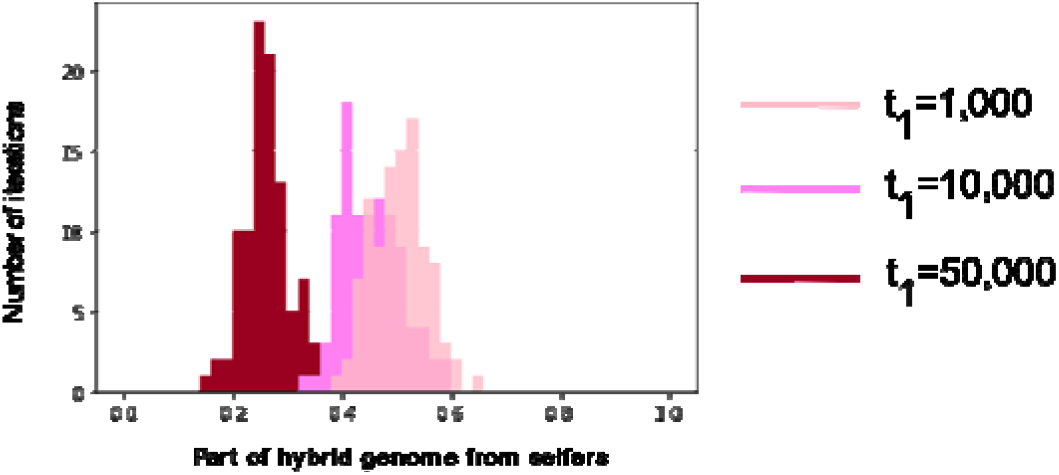
Hybrid genomic mixture depends on how much time passed before hybridization. We illustrate the histograms of the part of the hybrid genomes that were inherited from the self-fertilizing parental population for different number of generations of divergence between the parental populations prior to hybridization. We illustrate results for divergence times t_1_ = 1,000 (lighter pink), t_1_ = 10,000 (pink) and t_1_ = 50,000 (darker red).

Another effect of introducing some low rates of outcrossing in self-fertilizing hybrids is that it introduces recombination. This does not seem to have much of an effect when *r = 10^-6^*, as recombination rate is marginal. However, we see when *r = 10^-1^* that this recombination allows for the hybrid genome to exhibit mixed ancestry, instead of being composed of genes from only one or the other parental species.

### 3.4. Effects of divergence time on hybrid genome composition

The only case where hybrids’ genomes become a mix of genes from the two populations is when the hybrids outcross and recombination is frequent. In that case, we illustrate on Fig. 3 how the hybrid genome composition changes depending on the time of divergence prior to hybridization t_1_. As the populations have gone through longer divergence, the selferś genome is of worst quality compared with outcrosserś genomes (see Fig. 1). Consequently, and as illustrated on Fig. 3, the part of hybrid genome descending from self-fertilizing population ancestry tend to decrease with time prior to hybridization. Also, we see that the standard deviation of the self-fertilizing ancestry decreases: as differences between outcrossers and selfers amplify, results of hybridization are more stable from one iteration to the next.

## 4. Discussion

The transition from obligate outcrossing to predominantly selfing has happened multiple times in plants and animals (Jarne and Chalesworth 1993; Jarne and Auld 2006; The Tree of Sex Consortium 2014). This transition comes with a combination of phenotypic and genomic changes collectively known as the “selfing syndrome” (Shimizu and Tsuchimatsu 2015; Cutter 2019). Among those changes, outcrossers and selfers are expected to differ in their mutation load. Theory predicts that higher levels of heterozygosity in outcrossing species allows to better hide mutations that are simultaneously highly deleterious and highly recessive from selection, resulting in these reaching higher frequencies in outcrossers when compared to selfers (Wang *et al*. 1999). In contrast, theory also predicts that higher levels of homozygosity in self-fertilizing species result in less efficient recombination events, and thus stronger selective interference (Wright *et al*. 2013; Hartfield *et al*. 2017). As a result, slightly deleterious, co-dominant alleles are more likely to escape purging out by selection and increase in frequency by random genetic drift (Charlesworth *et al*. 1993). Our results generally agree with these predictions. As it turns out, there seems to be pros and cons for both outcrossing and self-fertilization in terms of mutation load. An important question is, which of the two processes is the strongest and most influences the genetic content of outcrossing and selfing species?

Our results, in line with other recent theoretical results (Sianta *et al*. 2022) emphasize the importance of the recombination rate in the build-up of the mutation load in outcrossers and selfers. We show that when recombination between adjacent loci is relatively high, outcrossers suffer little from selective interference. As a result, few deleterious mutations drift to fixation, fewer than in the self-fertilizing species that do suffer from selective interference. In this case, outcrossers evolve better genomes than their self-fertilizing counterparts (higher population mean fitness). Importantly, the prediction changes when we assume a low recombination rate between adjacent loci. In that case, outcrossers also suffer from selective interference since genetic associations are rarely re-shuffled. In addition, higher heterozygosity in outcrossers means higher frequency of deleterious, recessive mutations, as their detrimental effects are partially hidden in heterozygotes. These higher frequencies means that it is easier for these deleterious mutations to drift to fixation in outcrossers. As a result, we see that when recombination rates are low, outcrossers tend to accumulate more fixed deleterious mutations, which eventually translate into outcrossers evolving worse genomes (lower population mean fitness).

An interesting consequence of this result is that relative fitnesses may vary within genomes. Local rates of recombination are known to vary within genomes (Nachman and Churchill 1996; Kong *et al*. 2002; Jensen-Seaman *et al*. 2004). It is possible, depending on how other factors influencing selection and drift, that this could mean that areas of higher recombination rate exhibit higher mutation accumulation in a self-fertilizing species than in its outcrossing counterpart, while the reverse would hold true in areas of lower recombination rate.

Mutation accumulation by selective interference highly depends on genetic drift, and thus on the effective population size *N_e_.* On one hand, increasing population size limits the fixation of deleterious mutations. In simulations shown in Supplementary Material Figure S1, with *μ = 10^-5^* and *r = 10^-6^*, increasing population size from 500 to 5000 decreases the deleterious mutation fixation rate approximately by a factor 10. On the other hand, there are particular demographic events that leads to *N_e_* values much lower than the actual population size such as population bottlenecks or colonization events. Importantly, self-fertilizing species seem to be prone to this kind of events as they can in principle more easily colonize new environments and found new populations (Theologidis *et al*. 2014; NOEL *et al*. 2016; Grossenbacher *et al*. 2017). This may explain empirical patterns of relaxed purifying selection observed in some self-fertilizing species (SLOTTE *et al*. 2010; Burgarella *et al*. 2015; Wang *et al*. 2021). However, evidence for purifying selection in selfers is often mixed (Glémin *et al*. 2006; Haudry *et al*. 2008; Escobar *et al*. 2010). This mixed empirical evidence may be caused by the demographic history of the species as mentioned above, but also due to other factors such as divergence time between selfers and outcrossers, biased gene conversion, and stability of selfing as a reproductive strategy (Escobar *et al*. 2010). Overall, it is likely that self-fertilizing species suffer from greater mutation accumulation than what we predict here due to their demographic and evolutionary history.

How does mutation load in the parental species influence the ancestry of hybrids’ genes? Previous studies have hypothesized that hybrid individuals’ mating system should strongly impact the evolution of the genetic composition of hybrids and the levels of introgression to be expected in a hybrid zone. Pickup *et al*. (2019) predicts that in outcrossing hybrids, the many slightly deleterious, co-dominant mutations coming from the self-fertilizing ancestry should be selected against and purged while the highly deleterious, recessive mutations coming from the outcrossing ancestry should stay hidden in heterozygosity and be maintained. That is, the authors predict a purge of selfing ancestry in outcrossing hybrids. Similarly, they predict that in selfing hybrids, the highly deleterious, recessive mutations coming from the outcrossing ancestry should rapidly become homozygote and be selected against, while the many slightly deleterious, co-dominant mutations should keep on evading selection and be maintained. This time, the outcrossing ancestry is expected to be purged in selfing hybrids. Everything happens as if the mutation load was adapted to a certain reproductive mode and should be maintained in a hybrid that shares the same reproductive mode. This should result in limited introgression between outcrossing and self-fertilizing species, in both directions.

Though appealing, this verbal argument does not appear to be verified in our simulations. Our results emphasize that what matters is merely which parental species has the strongest mutation load. The genome ancestry with the worst fitness will most likely be purged in hybrids, independently of the hybrid mating system itself.

Kim *et al*. (2018) produced a similar model of introgression between outcrosser and selfer parental populations. In their simulations, they obtained that genomes from outcrossing populations are of better quality because selfer genomes accumulate fixed deleterious mutations by selective interference, predicting limited introgression of selfer into outcrosser genomes and greater introgression of outcrosser into selfer genomes. A few genomic parameters differ between our work and this paper. First, the authors use a gamma distribution for the selective coefficient of new deleterious mutations whose shape and scale parameters are such that overall selection is much weaker in their study. This may alter the mutation load balance by increasing genetic drift. However, it increases drift in both outcrossers and selfers, and simulations run with *s = 0.01* (which provides a selective coefficient distribution much closer to the one in Kim *et al*. (2018)) show an acceleration of deleterious mutation accumulation similar in the two parental species (see Supplementary Material Figure S2). Kim *et al*. (2018) also consider a genomic structure based on chromosome 1 of *Arabidopsis thaliana*. This notably means that they consider a greater number of loci under purifying selection, which is expected to increase the relative importance of selective interference and mutation accumulation. However, simulations with 1,000 and 10,000 instead of 100 loci (Supplementary Material Figure S3) again emphasize that mutation accumulation increases in both selfers and outcrossers; in both cases, outcrossers still accumulate more deleterious mutations in our simulations.

Focusing on the case of *Arabidopsis thaliana* Kim *et al*. (2018) acknowledge that they look at a scenario with relatively high recombination rates. As such, they may simply fall in a scenario where selective interference may play a stronger role in selfers (high recombination rates). How their result is generalizable to other self-fertilizing plant and animals remains to be assessed.

Mutation load is not the only factor that may influence introgression patterns between self-fertilizing and outcrossing species. Behavioral and physiological constraints are also major factors that are known to affect hybridization and introgression directions. In a hybrid zone between species with different mating systems, introgression is expected to happen more frequently from the selfing species to the outcrosser, than vice-versa (Kim *et al*. 2018; Pickup *et al*. 2019). This expectation relies on the premise that selfing (especially prior selfing (Tian-Bi *et al*. 2008; Brys *et al*. 2016; Berbel-Filho *et al*. 2021)) provides a very limited window of opportunity for outcrossing, by conspecific, heterospecific, or potential male hybrids (Pickup *et al*. 2019; Berbel-Filho *et al*. 2021). This pattern of higher levels of introgression from selfers into outcrossers has been commonly found in plant systems (Ruhsam *et al*. 2011; Ruhsam *et al*. 2013; Brandvain *et al*. 2014). Our findings extend that expectation, but from a different perspective. Our results indicate that under certain scenarios (here, lower recombination rates) self-fertilizing species may exhibit lower mutation load relative to outcrossing species. Following what we uncovered here, this should be expected to bias hybrid ancestry towards the selfer parents, that is to result in higher levels of introgression from selfers to outcrossers. However, under other scenarios (here, higher recombination rates), our model predicts the contrary: that hybrid ancestry would be biased towards the outcrossing, fitter parents.

We argue that our model provides insights into one of the factor, mutation load, that may influence the direction of introgression in hybrids zones between selfers and outcrossers. Of course, many other factors also influence such introgression and should be taken into account when comparing introgression patterns with parental species’ mutation load. For instance, the weak inbreeder/strong outbreeder hypothesis (WISO) (Brandvain and Haig 2005) states that due to the enhanced opportunity for genomic conflict, outcrossers’ gametes are more competitive than selfers’ ones. Given equal fertilization opportunities, if hybrids mostly outcross, higher introgression is expected from outcrossers to selfers under the WISO premises. Another factor that can contribute for deviations from the expectations of our model is selfing timing. Selfing can happen before (prior selfing), during (competitive), or after (delayed selfing) the opportunity for outcrossing (Lloyd 1979). The parental selfing time may limit the window of opportunity from an outcrossing hybrid to the selfing parental species, making the major direction of hybridization more common from selfer to outcrosser, than vice-versa (Brys *et al*. 2016; Berbel-Filho *et al*. 2021). Finally, unilateral genetic incompatibilities may bias the direction of introgression (Turelli and Moyle 2007) in both directions, regardless of the hybrids mating system and recombination rates. In our work, we focused solely on the effect of mating systems via mutation loads and have shown that parental species’ loads are good predictors of long-term genomic composition of hybrid species. Altogether, the examples presented above represent interesting avenues of research whenever deviations from this prediction are found.

## 5. Authors Contribution

FF and WMB-F designed the study. FF coded the model and ran the simulations. FF and WB-F interpreted the results and wrote the manuscript.

## 6. Acknowledgement

FF and WMB-F are supported by a NSFDEB-NERC Research Grant NE/T009322/1. We thank Christelle Fraisse, Benjamin C. Haller and Peter Ralph, along with editors and reviewers, for comments and suggestions that helped us improve the quality of the manuscript.

