## Supplementary Material for "Influence of the mutation load on the genomic composition of hybrids between outcrossing and self-fertilizing species"

### Mutation Load Build-Up in Parental Species

We present on Fig. S1 the characteristics of the mutational load born by two populations, one purely outcrossing and another one purely self-fertilizing.

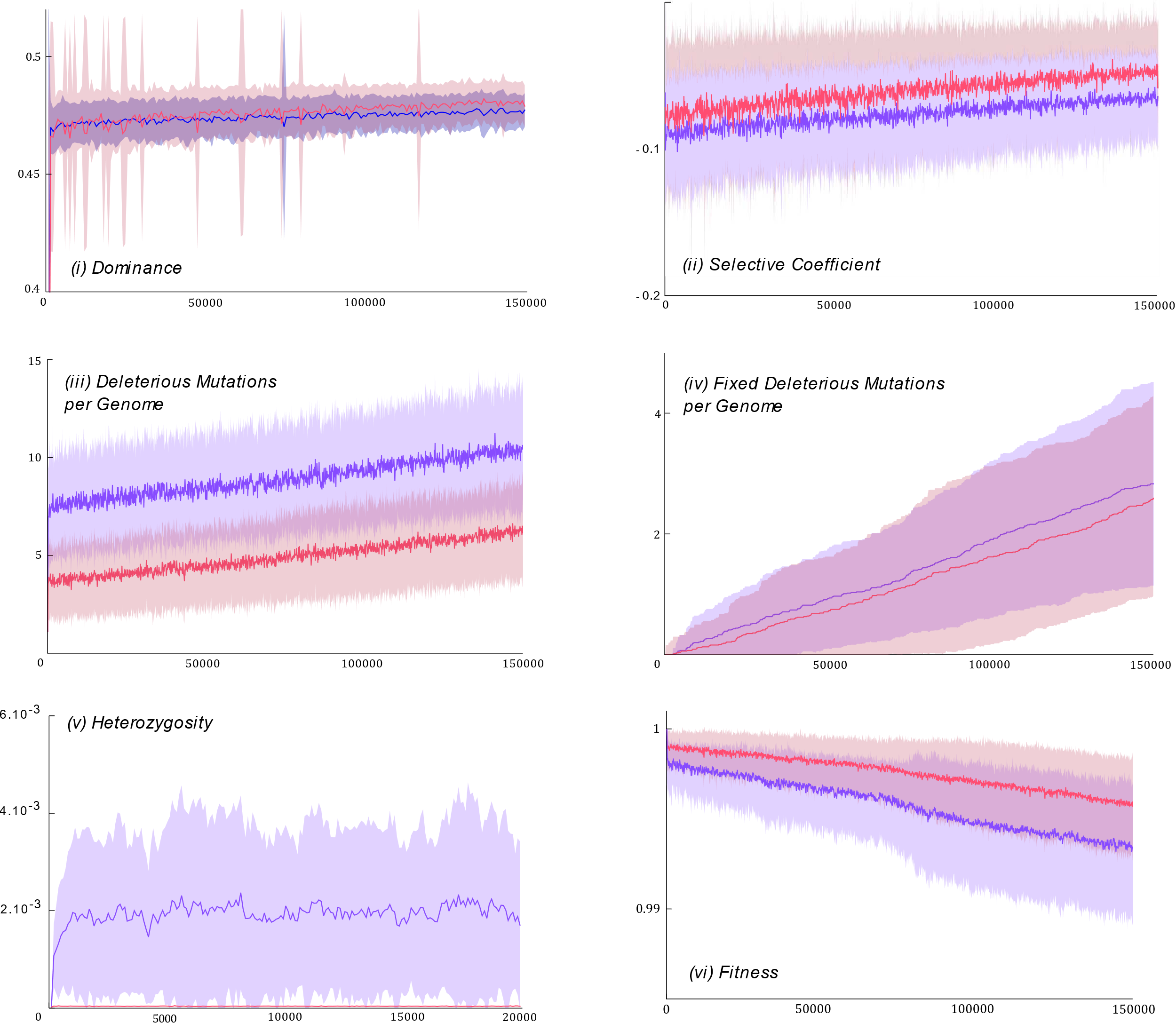

***Fig. S1****. Genomic composition of an outcrossing population (purple, selfing rate of σ_o_ = 0) and a self-fertilizing population (red, selfing rate of σ_s_ = 1) left to evolve during 150,000 generations (x-axis). Panels represent: (i) dominance of deleterious mutations, (ii) selective coefficients of deleterious mutations, (iii) number of deleterious mutations per genome, (iv) number of fixed deleterious mutations per genome and (v) heterozygosity and (vi) fitness of individuals. In each graph, plain lines represent mean values, while shaded areas represent standard deviation. Results are illustrated for the following parameter values: mutation rates μ = 10^-5^, recombination rate r = 10^-6^.*

The results found here are consistent with classic population genetics arguments. While outcrossers’ genomes bear a part of its polymorphism in a heterozygotic form, selfers’ genomes are mostly homozygous (Fig. S1.(i)). This leads to recessive mutations in selfers to be revealed to selection, in contrast with outcrossers, and thus to decrease in frequency or be eliminated altogether via purging. Because we assume a negative correlation between dominance of alleles and their detrimental effect on fitness, recessive mutations tend to be highly deleterious ones. Thus, selfers eliminate the highly deleterious mutations more than outcrossers, leading to the mean detrimental effect of deleterious mutations present in selfers’ genomes to be reduced in comparison with the mean detrimental effect of deleterious mutations present in outcrossers’ genome (Fig. S1.(ii)). High homozygosity in selfers strengthen the efficacy of selection, leading to a better purge of deleterious, recessive mutations and improving individuals’ fitness (Fig. S1.(v)). Because of this effect, less highly deleterious mutations are maintained as standing variation (polymorphism) in the self-fertilizing population (Fig. S1.(iii)). The total number of deleterious mutations per genome tends to increase with time. This is due to the progressive fixation of slightly deleterious mutations by genetic drift (the increase in Fig. S1.(iii) corresponds to the increase in Fig. S1.(iv)).

There does not seem to be a strong difference in the rates of random fixation of slightly deleterious alleles between the two populations. One mutation becomes fixed about every 55,000 generations in the outcrossing population, and about every 62,000 generations in the self-fertilizing population. Note that this difference is much smaller than the standard deviations of the two population. In accordance with classic population genetics results, the fixation rate in our simulations is of the order of the mutation rate (10^-5^).

Because the rate of fixation of deleterious mutations in selfers and outcrossers is similar, differences in selection and fitness are only due to the increase in selection efficacy in the self-fertilizing population associated with increased homozygosity. As a result, though the fitness of the two populations decrease at a steady, similar rate because of the fixation of deleterious mutations, the fitness of the selfers is consistently higher than the fitness of the outcrosser (Fig. S1.(v)). In the case illustrated here, the selfers have an overall better genome, a genome better purged from highly deleterious, recessive alleles.

We show on Figure S2 the accumulation of randomly fixed deleterious mutations for a diversity of parameter values.

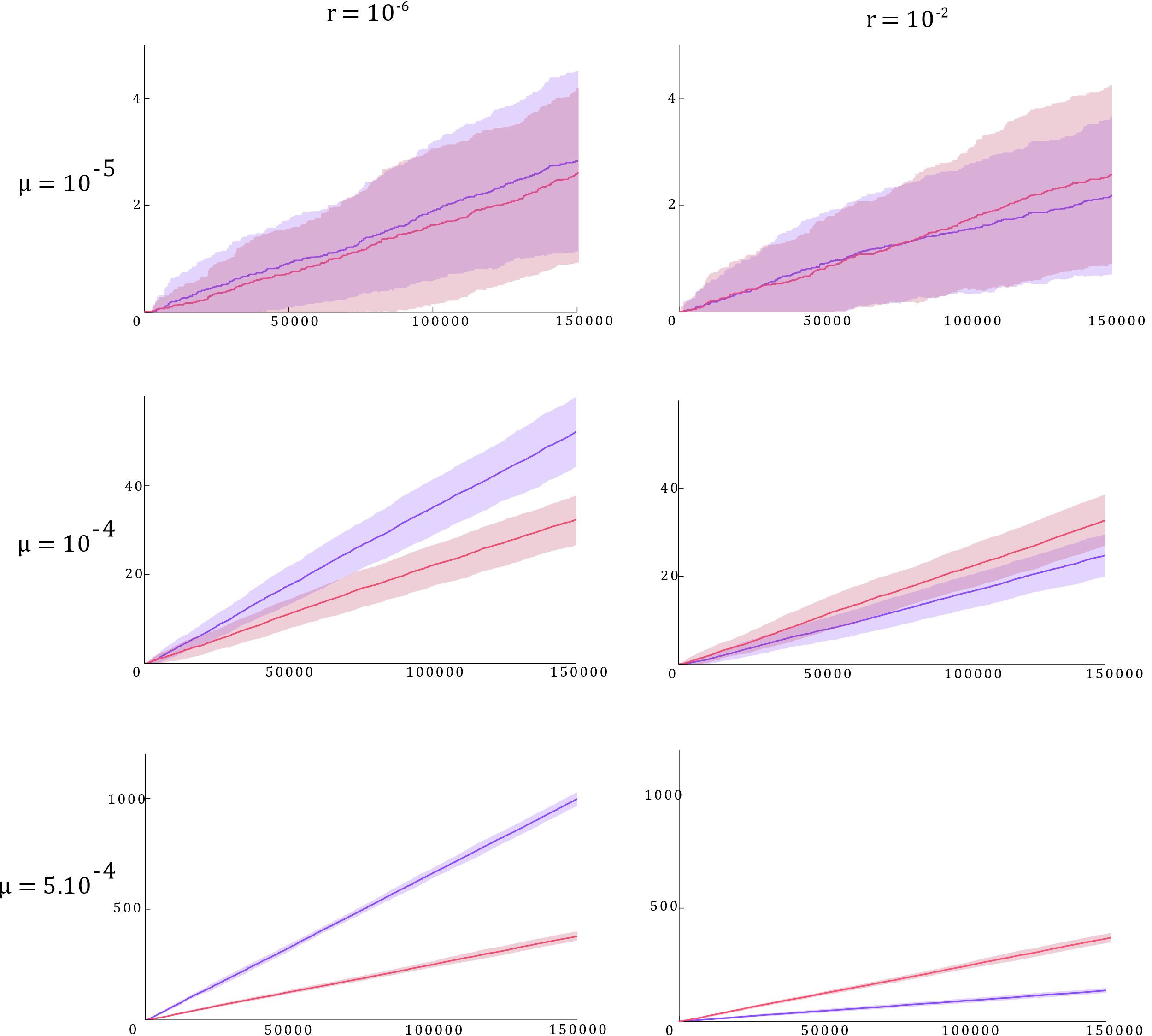

***Fig. S2.*** *Number of fixed deleterious mutations (y-axis) during 150,000 generations (x-axis) in an outcrossing population (purple, selfing rate of σ_o_ = 0) and a self-fertilizing population (red, selfing rate of σ_s_ = 1). In each graph, plain lines represent mean values, while shaded areas represent standard deviation. Values of r and μ are specified and change as columns and rows respectively.*

According to the relationship between the rate of random fixation of deleterious mutation and the mutation rate, we see the number of fixed mutations at any given time increasing with *μ*. Increased mutation rates, as it appears, make more obvious a trend that mostly depend on the recombination rate *r*. When *r* is low, both the outcrossing and the selfing populations have effectively low recombination rates. In that case, the outcrossing population appears to fix deleterious mutations more rapidly than the selfing population. This is because recessivity partly hides the detrimental effect of mutations when heterozygotes, which makes it easier for them to randomly drift from low to intermediate frequencies. Mutations that are more likely to reach intermediary frequencies are also more likely to drift to fixation. In contrast, when *r* is high, we see that the selfing population accumulates fixed mutations more rapidly than the outcrossing population. In fact, the rate of accumulation of fixed mutations in the selfing population does not change in comparison with the previous case of low *r* values – which was to be expected since recombination is ineffective in selfers and thus should not have any effect. The increase in *r* has led to a strong reduction in the accumulation of fixed deleterious mutations in the outcrossing population. This is because increased recombination rate improves selection efficacy in this population, leading to a decrease of the probability that a given deleterious mutation can drift to fixation by chance. Here, we see two antagonistic effects. First, selection efficacy is reduced in outcrossers due to heterozygosity and recessivity. Second, selection efficacy is increased in outcrosser due to recombination and decreased selection interference. We note that the second effect can overcome the first only if (i) the mutation rate or the evolutionary time considered are large; and if (ii) recombination in the outcrossing population is high enough in the genomic segment considered so that it prevents much of selective interference effects.

### Alternative Parameter Values

In the above simulations, we used population sizes of *N_pop_ =* *500* individuals, a mean selective effect of deleterious mutations of *1/λ = 0.1*, and a number of loci under selection of *N_loc_ = 100*. These have been chosen to limit the computational power needed to run the simulations. However, they do not correspond necessarily to realistic values. In the following, we illustrate results for alternative parameter values.

#### *Npop*

We illustrate on Fig S3 the effect of increasing population sizes. We see that increasing population sizes by a factor 10 reduces accumulation of deleterious mutations also by a factor 10, which is to be expected. As population size increases, fixation of deleterious mutations by drift is less probable. Importantly, we see that the self-fertilizing and the outcrossing populations’ accumulation of deleterious mutations are reduced very similarly – population size does not appear to impact the balance of deleterious mutations accumulation between the two populations.

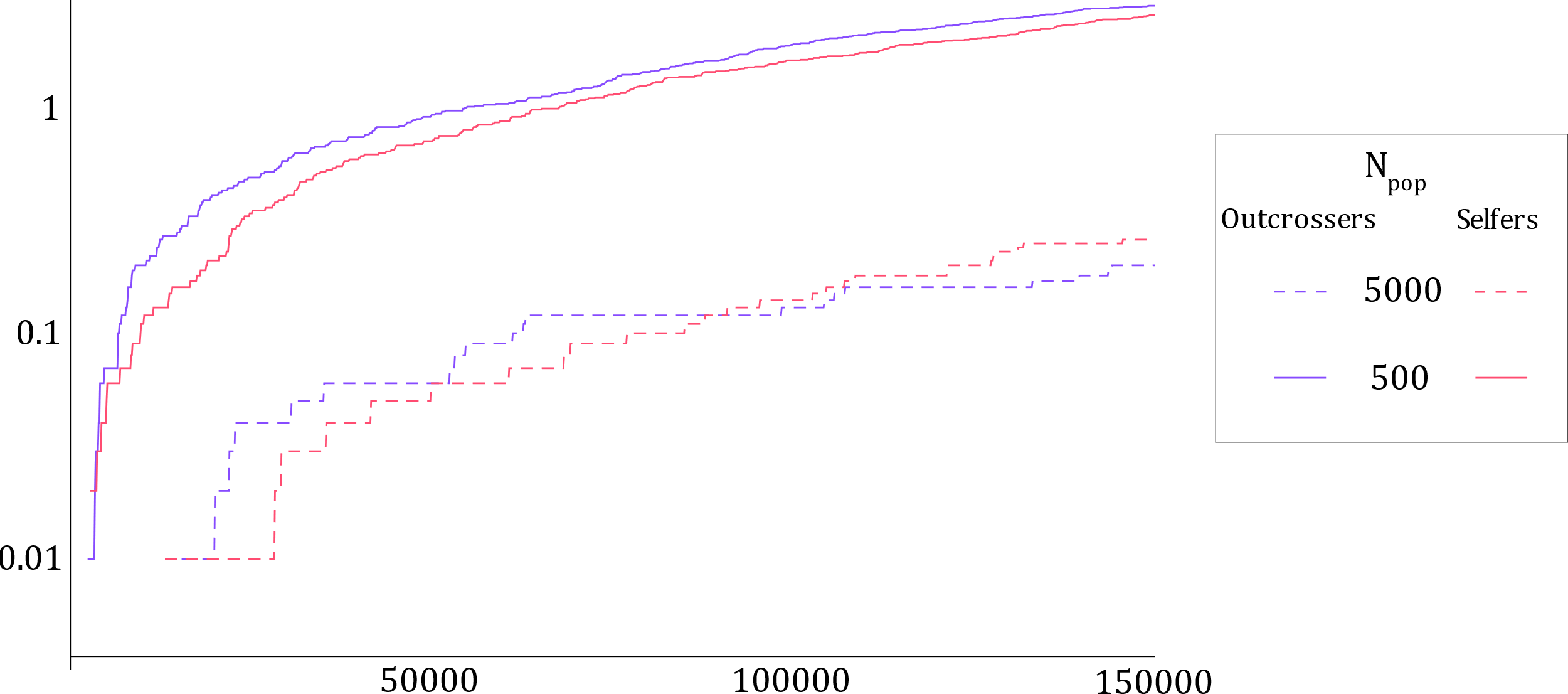

***Fig. S3.*** *Average (out of 100 iterations) accumulation of deleterious mutations (y-axis) during 150,000 generations (x-axis) in purely outcrossing (purple) and self-fertilizing populations (red). Two values of population size N_pop_ are illustrated: 500 (as in the main text, plain lines) and 5,000 (dashed lines).*

## *1/λ*

We illustrate on Fig S4 the effect of decreasing by a factor 10 the mean detrimental effect of mutations. The value of *1/λ = 0.01* has been chosen as it allows to produce a distribution of deleterious mutations’ effects similar to Kim et al., 2018. We see that this tends to accelerate the fixation of deleterious mutations, as it reduces the selection against them. Again, it is important to note that the accumulation of deleterious mutations of both populations is increased similarly.

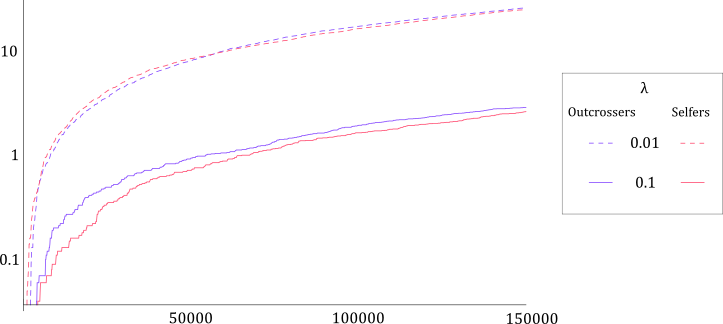

***Fig. S4.*** *Average (out of 100 iterations) accumulation of deleterious mutations (y-axis) during 150,000 generations (x-axis) in purely outcrossing (purple) and self-fertilizing populations (red). Two values of selection parameter λ are illustrated: 0.1 (as in the main text, plain lines) and 0.01 (similar to Kim et al., 2018, dashed lines).*

#### *N_loc_*

We show on Fig. S5 the effect of increasing the number of loci under selection in the genomes. It appears that increasing *N_loc_* leads to an increase of deleterious mutation accumulation. This is due to an increase in selective interference facilitating the fixation of deleterious mutations. The increase in mutation accumulation appears to be proportional to *N_loc_*, and again similar in the two populations.

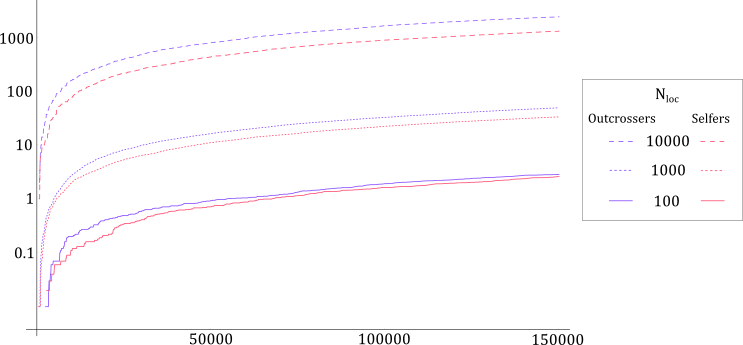

***Fig. S5.*** *Average (out of 100 iterations) accumulation of deleterious mutations (y-axis) during 150,000 generations (x-axis) in purely outcrossing (purple) and self-fertilizing populations (red). Three values of locus number N_loc_ are illustrated: 100 (as in the main text, plain lines), 1,000 (dotted lines) and 10,000 (dashed lines).*
